# GPT-4 based AI agents – the new expert system for detection of antimicrobial resistance mechanisms?

**DOI:** 10.1101/2024.05.06.592800

**Authors:** Christian G. Giske, Michelle Bressan, Farah Fiechter, Vladimira Hinic, Stefano Mancini, Oliver Nolte, Adrian Egli

**Author notes:** **Correspondence:** Prof. Adrian Egli, MD PhD, Institute of Medical Microbiology, University of Zurich, Gloriastrasse 30, 8006 Zurich. equal contribution (Oliver Nolte, Adrian Egli). equal contribution (Christian G. Giske, Michelle Bressan).

## Abstract

**Background:** EUCAST recommends a two-step process for beta-lactamases in Gram-negative bacteria. Screening with minimal inhibitory concentrations (MICs) or inhibition zone diameters for potential extended spectrum beta-lactamase (ESBL), plasmid-mediated AmpC beta-lactamase, or carbapenemase production is followed by confirmatory tests. GPT-4 and its newly released customized GPT-agent may support the initial EUCAST-screening process. We aimed to validate a customized GPT-agent to identify potential resistance mechanisms.

**Methods:** We used 225 Gram-negative isolates. Based on phenotypic resistances against beta-lactam antibiotics, we formed four categories: “none”, “ESBL”, “AmpC”, or “carbapenemase”. We included 862 phenotypic categories. Next, we customized a GPT-agent with EUCAST-guidelines, expert rules, and EUCAST-breakpoint table (v13.1). We compared routine diagnostic outputs (reference) to (i) EUCAST-GPT-expert, (ii) medical microbiologists, and (iii) GPT-4 without customization. We determined performance as sensitivities and specificities to flag suspect resistance mechanisms.

**Results:** Three human readers showed concordance in 814/862 (94.4%) phenotypic categories and used in median eight words (IQR 4-11) for reasoning. Median sensitivity and specificity for ESBL, AmpC, and carbapenemase were 98%/99.1%, 96.8%/97.1%, and 95.5%/98.5%, respectively. Three independent prompting rounds of the GPT-agent showed concordance in 706/862 (81.9%) categories but used in median 158 words (IQR 140-174) for reasoning,. Median sensitivity and specificity for ESBL, AmpC, and carbapenemase prediction were 95.4%/69.23%, 96.9%/86.3%, and 100%/98.8%, respectively. In the non-customized GPT-4, 169/862 (19.6%) categories could be interpreted. Of these 137/169 (81.1%) categories agreed with routine diagnostic. The non-customized GPT-4 used in median 85 words (IQR 72-105) for reasoning.

**Conclusion:** Human experts showed higher concordance and shorter argumentations compared to GPT-agents. Human experts showed comparable median sensitivities and higher specificities compared to GPT-agents. GPT-agents showed more unspecific flagging of ESBL and AmpC, potentially, resulting in additional testing, diagnostic delays, and higher costs. GPT-4 and GPT-agents are not IVDR/FDA-approved, but validation of LLMs is critical and datasets for benchmarking are needed.

## Introduction

As the healthcare sector grapples with the escalating challenge of antimicrobial resistance (AMR), the need for advanced diagnostic methods increases (1). For beta-lactamases in Gram-negative bacteria this is usually a two-step process (2): first, based on screening breakpoints for minimal inhibitory concentrations (MICs) or inhibition zone diameters suspected isolates with potential extended spectrum beta-lactamase (ESBL), plasmid-mediated AmpC, and carbapenemase production are flagged; second, suspected resistance is then confirmed with additional tests *e*.*g*., molecular assay for specific resistance genes (3).

Kirby-Bauer disk diffusion is a commonly used method for determining bacterial susceptibility and exemplifies the complexities of microbiological diagnostics (4, 5). This is particularly true for detection of extended-spectrum beta-lactamases (ESBLs), AmpC beta-lactamases, and carbapenemases. Precise interpretation within this framework is vital, yet challenged by technical and human variability, and the necessity for continual expert knowledge updating. Reproducibility in reading and interpretation of disk diffusion has been reported to be variable (6-8).

In this context, the potential integration of AI technologies, such as generative models like GPT-4, customized GPT-agents (openAI), or other large language models (LLMs) into laboratory medicine is an area of growing interest (9). However, it is crucial to note that AI-tools are not routinely established due to the current absence of compliance with In Vitro Diagnostic Regulation (IVDR) and Food and Drug Administration (FDA) regulations (10). This regulatory gap underscores the importance of validation to ensure their reliability, accuracy, and safety in clinical diagnostics.

Our study aimed to contribute to this validation process. By utilizing GPT-4 and a customized GPT-agent to interpret test results. We aimed to understand how such AI-tools can be calibrated and utilized within the stringent frameworks of laboratory medicine. The study provides an opportunity to gather valuable insights into the integration of AI in diagnostics, for future validation and regulatory approval. This is a critical step in ensuring that AI-tools can be safely and effectively used to enhance patient care in clinical laboratories (9, 11, 12).

## Methods

### Study Design and Sample Collection

We conducted a retrospective study involving 225 Gram-negative isolates from routine diagnostics. Laboratory processes are ISO/IEC accredited. We randomly included four *Acinetobacter baumannii*, three *Citrobacter freundii*, two *C. koseri*, 13 *Enterobacter cloacae* complex, 132 *Escherichia coli*, five *Klebsiella aerogenes*, five *K. oxytoca*, 40 *K. pneumoniae*, three *Morganella morganii*, ten *Proteus mirabilis*, one *P. vulgaris*, one *Pseudomonas aeruginosa*, and six *Serratia marcescens*. The isolates measured with disk diffusion according to EUCAST-guidelines (13) using antibiotic discs purchased from i2a (Perols Cedex, France) and Mueller-Hinton agar plates (BD, Franklin Lakes, NJ). The SIRweb/SIRscan system (i2a) was used to measure the inhibition zone diameters (8). The automated SIRweb expert system based on EUCAST-guidelines for the detection of resistance mechanisms enabled the categorization of four potential phenotypic resistance categories: suspected «none» (n=75), «ESBL» (n=111), «AmpC» (n=32), and «carbapenemase» (n=23). A total of 13.8% of isolates were excluded. Main reasons were poor image quality, non-resulting interpretation by GPT-4, or no valid interpretation guideline e.g., in *A. baumannii* and *P. aeruginosa* for AmpC. We finally included a total of 862 valid category phenotypes for subsequent analysis. All data can be downloaded at (**weblink upon publication**).

### Ethics and data protection

The study focused only on quality assessment of a new diagnostics tool and was not including any patient characteristics. No personal or patient-related data was shared with GPT-4 or the customized GPT-agent. The study was approved by the local ethical committee (Req-2023-00752).

### Development of the Customized GPT-Agent

We generated a customized GPT-agent named “EUCAST-GPT-expert” using the commercial version of ChatGPT (openAI). The specific GPT-4 version used was dated 06/11/2023 to assist in the interpretation of EUCAST antimicrobial susceptibility testing. The GPT-agent was customized through the following steps (**Figure 1**):

**Figure 1.**
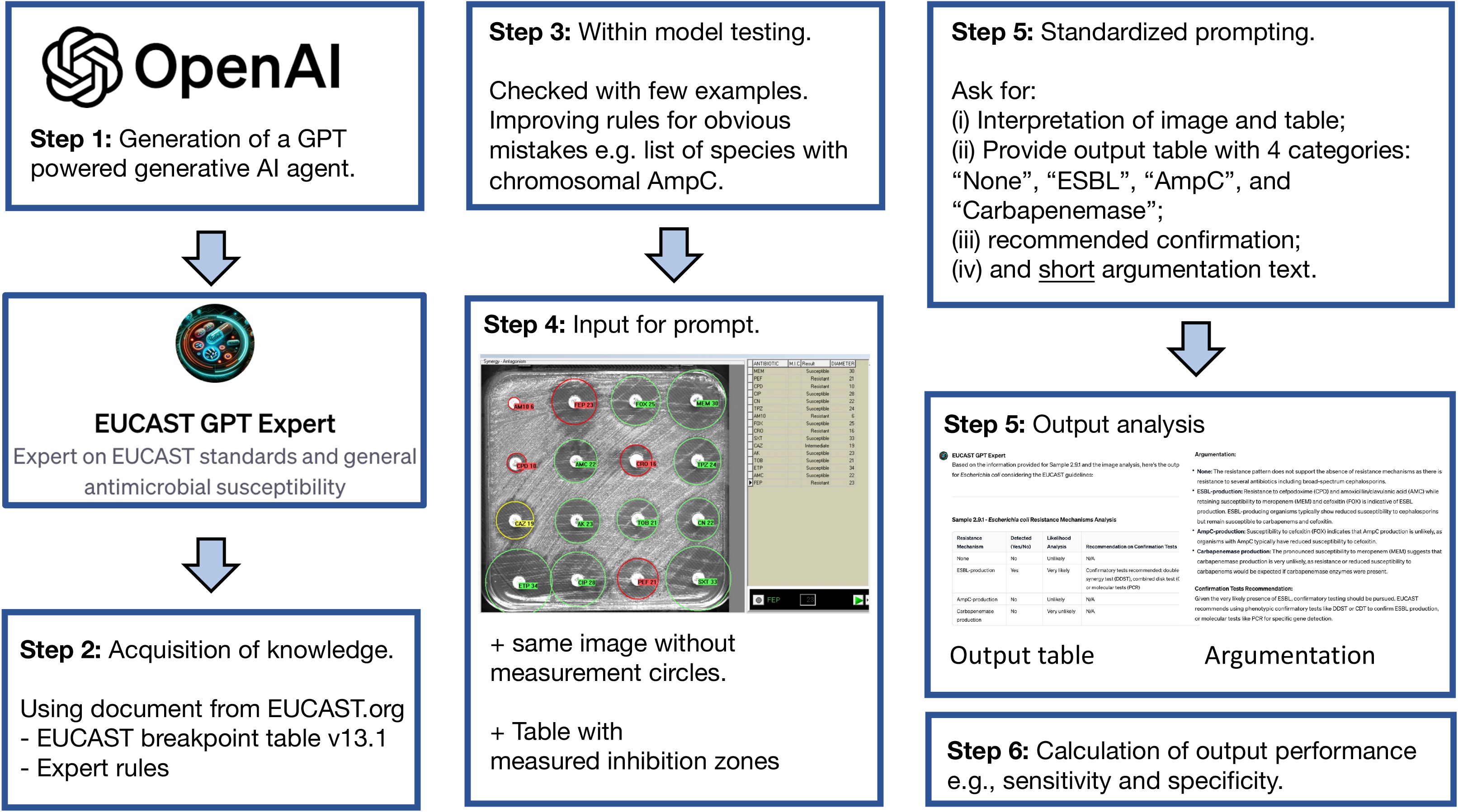
Workflow for validation of GPT-4 based generative AI-agent.

1. *Knowledge Acquisition*. The GPT-agent was equipped with the following documents: latest breakpoint tables (v13.1, (14)), EUCAST-guidelines for detection of resistance mechanisms and specific resistances of clinical and/or epidemiological importance (v2, July 2017, (3)), and EUCAST expert rules (15, 16). This knowledge base was intended to enable the GPT-agent to understand and apply EUCAST-guidelines accurately.
2. *Model Refinement*. Preliminary testing was performed with some examples to fine-tune its interpretative capabilities. Corrections were made to ensure the GPT-agent did not repeat identifiable errors, such as miss-listing species known to have chromosomal AmpC with common de-repression, e.g., *Citrobacter freundii, Enterobacter cloacae, Klebsiella aerogenes*, etc. or providing awareness of intrinsic resistances such as *a*mpicillin in *Klebsiella* spp..
3. *Input Preparation*. For each isolate, the input comprised images of disk diffusion plates from SIRscan. Accompanying these images was a table detailing the measured inhibition zones for each antibiotic (**Supplementary Figure 1 and 2, Supplementary Table 1**).
4. *Standardized Prompting*. The GPT-agent was prompted in a structured manner to interpret the provided data (see below for exact prompt). The same prompt was also used for the non-customized GPT-4 and provided to human experts. Each group was tasked with categorizing the resistance level into one of four categories and to elaborate on its reasoning in a brief argumentative text.
5. *Output Analysis*. The analysis included a detailed output table containing the resistance categories and a section where the GPT-agent provided its argumentation for each categorization. For each bacterial isolate, the agent was prompted to analyse images and generate an output table identifying the resistance mechanisms. The agent was instructed to force a binary (yes/no) decision on the presence of resistance mechanisms and to assess the likelihood of each identified mechanism.

### Standardized Prompting Procedure

For each sample, the identical queries were used. Example “Sample 6.70.1. Escherichia coli - Make an output table. In that output table identify the resistance mechanisms you have detected from the analysis of the provided images and information: (i) None, (ii) ESBL-production, (iii) AmpC-production, or (iv) Carbapenemase production. Make a specific call with yes/no answers - force yourself to provide an answer. Add into this table a likelihood analysis for each resistance mechanism: (a) very likely, (b) likely, (c) unlikely, (d) very unlikely. For samples with likely or very likely results make a recommendation on the potential confirmation tests which should be used according to EUCAST. In the title of each table mention the sample ID and the bacterial species. Provide a short argumentation for each resistance phenotypes (none, ESBL, AmpC, and Carbapenemase) based on the measurement and image analysis.”

### Output Analysis and Argumentation

The output table from the EUCAST-GPT-expert included sample identification and resistance mechanism detection, with a likelihood analysis (**Supplementary Table 2**). The GPT-agent was also required to provide a short argumentation for each decision, drawing on the measured inhibition zones and image analysis. This approach aimed to mimic the reasoning process of human experts.

### Benchmarking and Validation

The outputs of all groups, GPT-4, GPT-agent, and three medical microbiologists were compared against the previously reported results in routine diagnostics (reference standard). As there was some variability in the interpretation of the results, we showed the median outputs of the three microbiological experts and performed three independent prompting rounds for the EUCAST-GPT-expert, where also the median outputs were used. The output of the non-customized GPT-4 was poor, and therefore we did not repeat the prompting. We calculated concordance rates for categories identified by human experts and the EUCAST-GPT-expert. We recorded the median number of words used for reasoning.

### Statistical Analysis

Descriptive statistics were used to summarize the performance of the human experts, the EUCAST-GPT-expert and non-customized GPT-4. Sensitivity, specificity, and negative and positive predictive value were compared. The median sensitivities and specificities for human readers was calculated with interquartile ranges (https://www.medcalc.org) and compared to evaluate the diagnostic accuracy of the AI-models.

## Results

The customized GPT-agent (“EUCAST-GPT-expert”), informed by EUCAST-guidelines and trained on expert rules, analyzed 862 phenotypes from 225 Gram-negative bacterial isolates. When compared to the reference standards, the GPT-agent demonstrated a median sensitivity of 95.4% for suspected ESBL detection, 96.9% for suspected AmpC, and 100% for suspected carbapenemase detection. Specificity, however, varied, with 69.2% for ESBL, 86.3% for AmpC, and 98.8% for carbapenemases, indicating a propensity for over-flagging potential resistances, particularly for ESBL and AmpC (**Table 1**). The GPT-agent’s tendency to over-flag, particularly ESBL-producing isolates with amoxicillin resistance and amoxicillin/clavulanic acid susceptibility due to a narrow spectrum beta-lactamase, might lead to unnecessary confirmatory testing and higher costs. Similarly, the over-flagging of AmpC with cefoxitin susceptibility suggests areas for model refinement. Of note, when the GPT-agent was asked to re-analyze and carefully consider the knowledge and rules, then the output was often adapted and corrected. For this analysis, we have however used only the first reply of the GPT-agent.

**Table 1.**
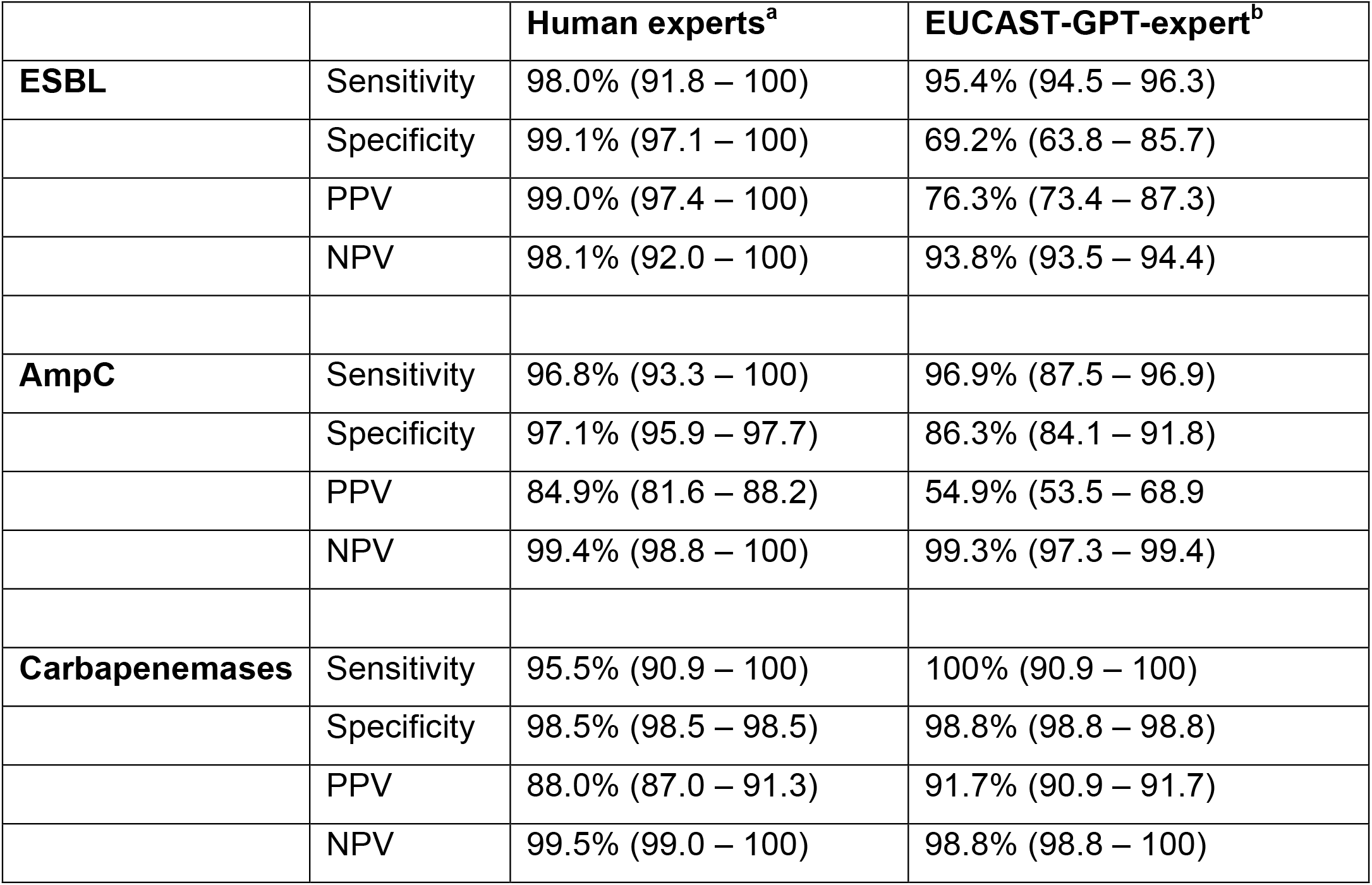
Sensitivity and specificity of human experts and the customized EUCAST-GPT-expert. ^a^, three human experts (median). ^b^, three independent prompting outputs from the customized GPT-4 agent <EUCAST-GPT-expert=. As reference standard, we used the results reported according to our ISO-accredited laboratory information system. ESBL, extended spectrum beta-lactamase; None, no specific molecular resistance mechanism.

Interestingly, when we focused on individual bacterial species, we observed a potential species-specific effect. As an example, in *E. coli* (n=132) we noted lower ESBL detection rates compared to all samples (median sensitivity 86.4% vs. 95.4%), and a lower median sensitivity compared to *K. pneumoniae* and *K. oxytoca* (n=45, median sensitivity 100%, **Table 2**). However, the specificity in ESBL-producing *E. coli* was higher compared to all samples (median specificity 76.9% vs. 69.2%), and higher compared to *K. pneumoniae* and *K. oxytoca* (median specificity 61.9%). This could potentially be explained by the previously mentioned misinterpretation of ESBL in the case of amoxicillin resistance and amoxicillin/clavulanic acid susceptibility.

**Table 2.**
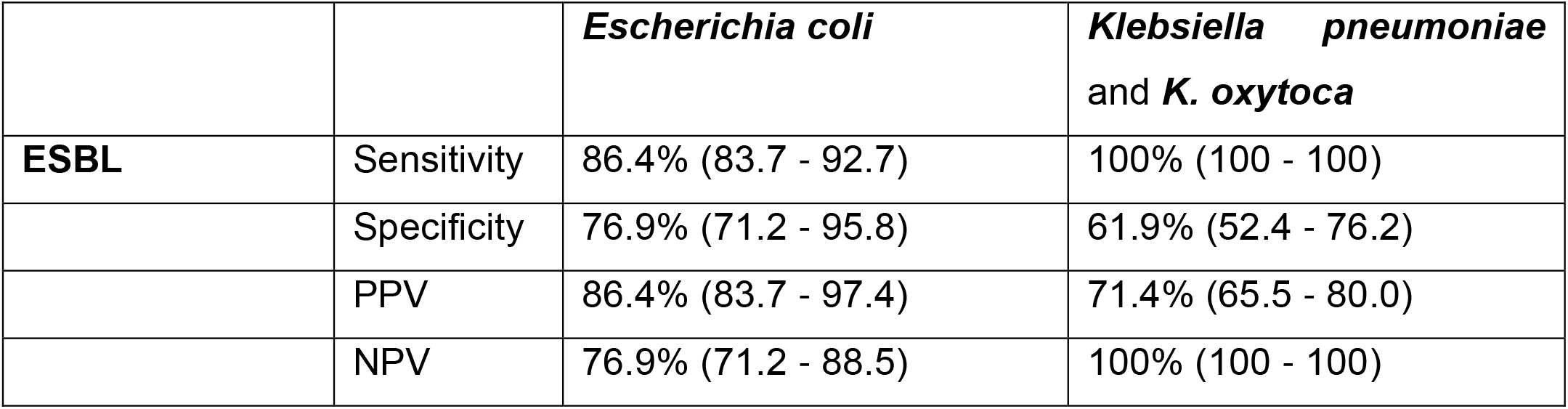
Comparison of common bacterial species and the performance of the EUCAST-GPT-expert. Only ESBL was analyzed, as for AmpC and carbapenemase producing bacteria the numbers where too low and not balanced. Outputs of K. pneumoniae and K. oxytoca were pool to generate more robust numbers.

|  |  | *Escherichia coli* | *Klebsiella pneumoniae*<br>and *K. oxytoca* |
| --- | --- | --- | --- |
| **ESBL** | Sensitivity | 86.4% (83.7 - 92.7) | 100% (100 - 100) |
|  | Specificity | 76.9% (71.2 - 95.8) | 61.9% (52.4 - 76.2) |
|  | PPV | 86.4% (83.7 - 97.4) | 71.4% (65.5 - 80.0) |
|  | NPV | 76.9% (71.2 - 88.5) | 100% (100 - 100) |

### Comparison of AI and Human Performance

Human experts generally exhibited higher specificity, particularly in the detection of ESBL and AmpC phenotypes, compared to the EUCAST-GPT-expert. However, the sensitivity was comparable between the human experts and the AI-agent (**Table 1**). Positive and negative predictive values were in general also higher in human experts compared to EUCAST-GPT-expert (**Table 1**). We also wanted to explore the performance of a customized GPT-agent compared to a non-customized GPT-4 prompt. Therefore, we also generated outputs using the same images and prompting strategy but with a non-customized GPT-4. In the non-customized GPT-4, only 169/862 (19.6%) categories could be interpreted. Of these 137/169 (81.1%) categories agreed with routine diagnostic. For this subgroup, the available phenotypic categories were too low to provide robust enough calculations for sensitivities and specificities for individual resistance mechanisms.

### Analysis of Argumentation

The EUCAST-GPT-Expert’s argumentation was more detailed than replies from human experts, which could be beneficial for educational purposes or in complex cases where thorough explanations are warranted. However, this verbosity may not be practical in a routine diagnostic setting where brevity is preferred. The human experts used in median eight words (IQR 4-11), indicating concise rationales for their decisions. In contrast, the EUCAST-GPT-expert provided more extensive explanations, with a median of 158 words (IQR 140-174) and suggested in addition also confirmation steps with a median of five words (IQR 4-9). Although we did not perform an in-depth analysis of the quality, we noted that the EUCAST-GPT-expert provided in some cases correct interpretations of *e*.*g*. the phenotypic resistance category, but the argumentation for the interpretation was not correct.

EUCAST-GPT-expert provided an additional text to specifically recommend a next confirmational step. In most situations a correct follow-up assay was described, *e*.*g*., with a specific PCR, but occasionally it was simply noted that a confirmational assay is necessary.

The non-customized GPT-4 used in median 85 words (IQR 72-105) for reasoning and 0 words (IQR 0-0) to suggest confirmation steps. In very few situations the next confirmational steps were properly described.

### Human Expert and EUCAST-GPT-expert concordance

The three human experts showed in 814/862 (94.4%) concordance across all phenotypic categories. Concordance for ESBL, AmpC, and carbapenemases were 94.0%, 96.6%, and 98.6%, respectively. In contrast, three separated runs with the EUCAST-GPT-expert showed in 706/862 (81.9%) phenotypic concordance across the categories. Concordance for ESBL, AmpC, and carbapenemase was 74.4%, 72.3%, and 97.7%, respectively.

## Discussion

We highlight the potential and limitations of integrating a customized AI-agent, specifically GPT-4 and GPT-agents, in interpreting antimicrobial susceptibility tests. The GPT-agent’s comparable sensitivity to human experts in detecting ESBL, AmpC beta-lactamases, and carbapenemases is promising and underscores its ability to identify resistance mechanisms effectively, a crucial aspect in combating AMR. However, the lower specificity and over-flag of certain resistances call for a careful approach during integration in clinical practice as it may result in critical delays as well as increased workloads and costs during confirmation. The detailed argumentation provided by the GPT-agent, while beneficial for educational purposes, may need to be streamlined for practical application in busy diagnostic labs. The customized-GPT agent showed a clear benefit over a non-customized version.

The variability in interpretation among human experts illustrates the subjective nature of manual readings and the potential for AI to provide more standardized interpretations (8). On this occasion also different interpretation of AmpC, considering only plasmid-mediated or both plasmid- and chromosomal-mediated resistance. Variability in plate reading and interpretation, and as a matter of fact also human error needs to be considered in any diagnostic test validation. However, the expertise and clinical judgment of human microbiologists remain invaluable, especially in complex or ambiguous cases. Also different LLM models may show variable results as indicated in a comparison of clinical microbiology scenarios between GPT-3.5 and GPT-4 (17). Importantly, GPT-4 and the customized-GPT agent do not provide detailed insights on how the data is analysed and interpreted. These systems remain a black box. Open-source LLMs, such as LLAMA-2, will become very important to explore the technology and understand how AI algorithms work with real-world data (9).

Our results demonstrate the importance of ongoing AI refinement, considering both the rapid evolution of AMR patterns and advancements in technology. The study also underscores the necessity of regulatory compliance and validation for AI-tools in healthcare, as highlighted by the lack of IVDR/FDA approval for GPT-4 in clinical diagnostics (9). Thereby, our study may also be seen as a blueprint for a dataset which allows tracking of LLM progress and as a benchmark *in silico* dataset. Repetitive testing of the same dataset allows tracking performance evolution of LLMs.

Future studies should focus on expanding the dataset to include a wider range of bacterial species and resistance mechanisms. Additionally, exploring AI’s role in interpreting other diagnostic tests could provide a more comprehensive understanding of its capabilities and limitations (11). Collaborations between AI developers, medical microbiologists, and regulatory bodies are essential to ensure that AI-tools are safely and effectively integrated into clinical workflows.

Our study has important limitations. It is limited by the specific version of GPT-4 used, as this field rapidly evolves and releases are published on a regular basis, by the time this article is published, the performance has likely improved. Also, we have focused on Gram-negative bacteria and beta-lactamases. Future studies should explore different bacterial species and resistance mechanisms e.g. with methicillin-resistant *Staphylococcus aureus* or vancomycin-resistant *Enterococcus faecium*. Moreover, the AI’s performance in a real-world clinical setting may differ from this controlled study environment. Prospective trials are needed in the field of AI and laboratory investigations; however, a first step must be a retrospective assessment to ensure its safety and baseline performance. Next, we have only few *Acinetobacter baumannii* and *Pseudomonas aeruginosa* isolates included, future work needs a more balanced dataset. Finally, our dataset has only been used in a GPT-4 and GPT-agent related context and not been used to explore other LLMs.

## Acknowledgments

We want to thank the laboratory technical personnel at the Institute of Medical Microbiology at the University of Zurich for processing and measurements of antimicrobial resistance profiles.

## Supplementary material

**Supplementary Figure 1.**
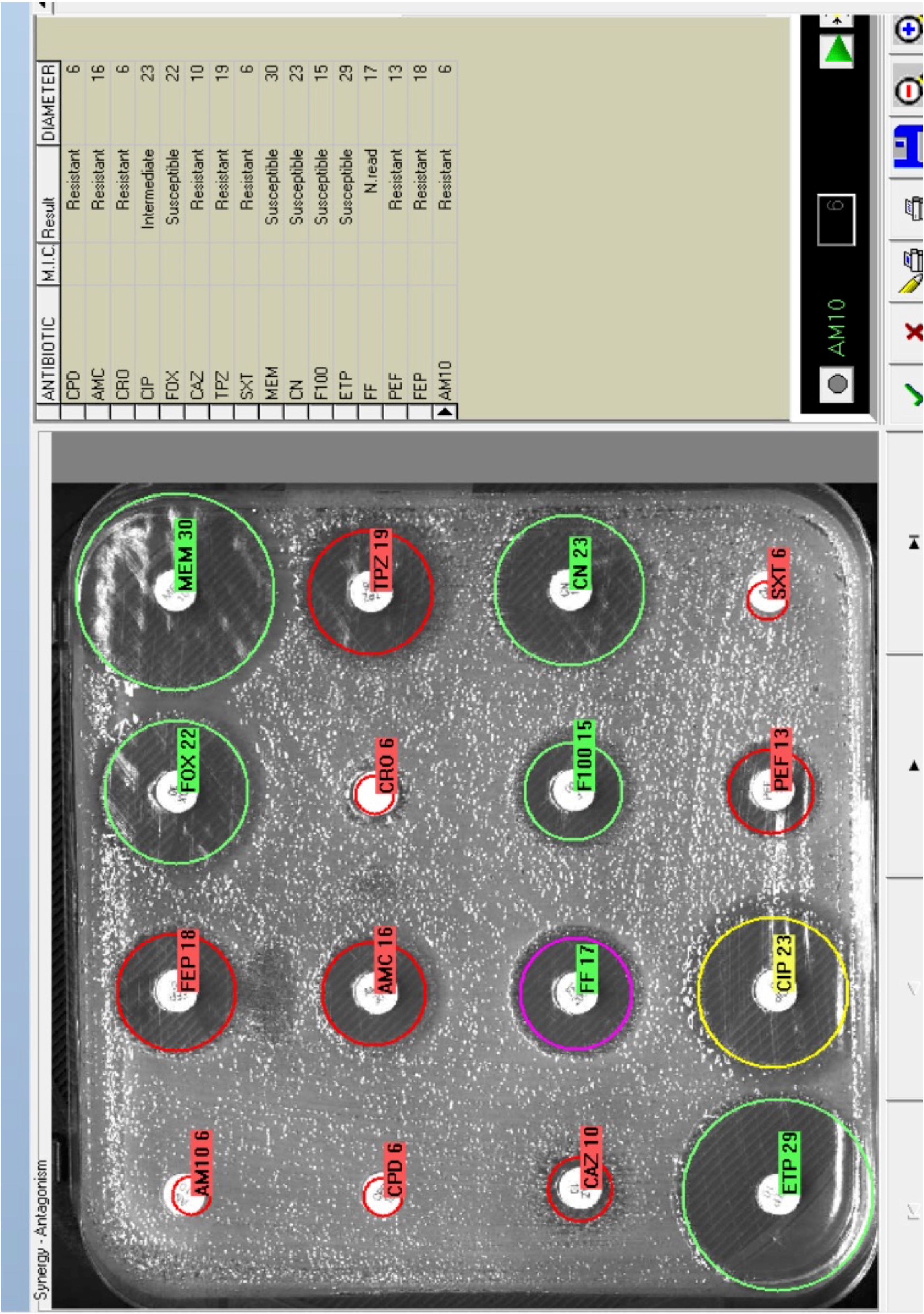
Representative photography of a disk diffusion assay generated by the SIRscan device. The image originates from routine diagnostics and shows measured circles to indicate the inhibition zone diameter in mm and a categorical interpretation according to EUCAST. As representative example image 2.3.1 was used.

**Supplementary Figure 2.**
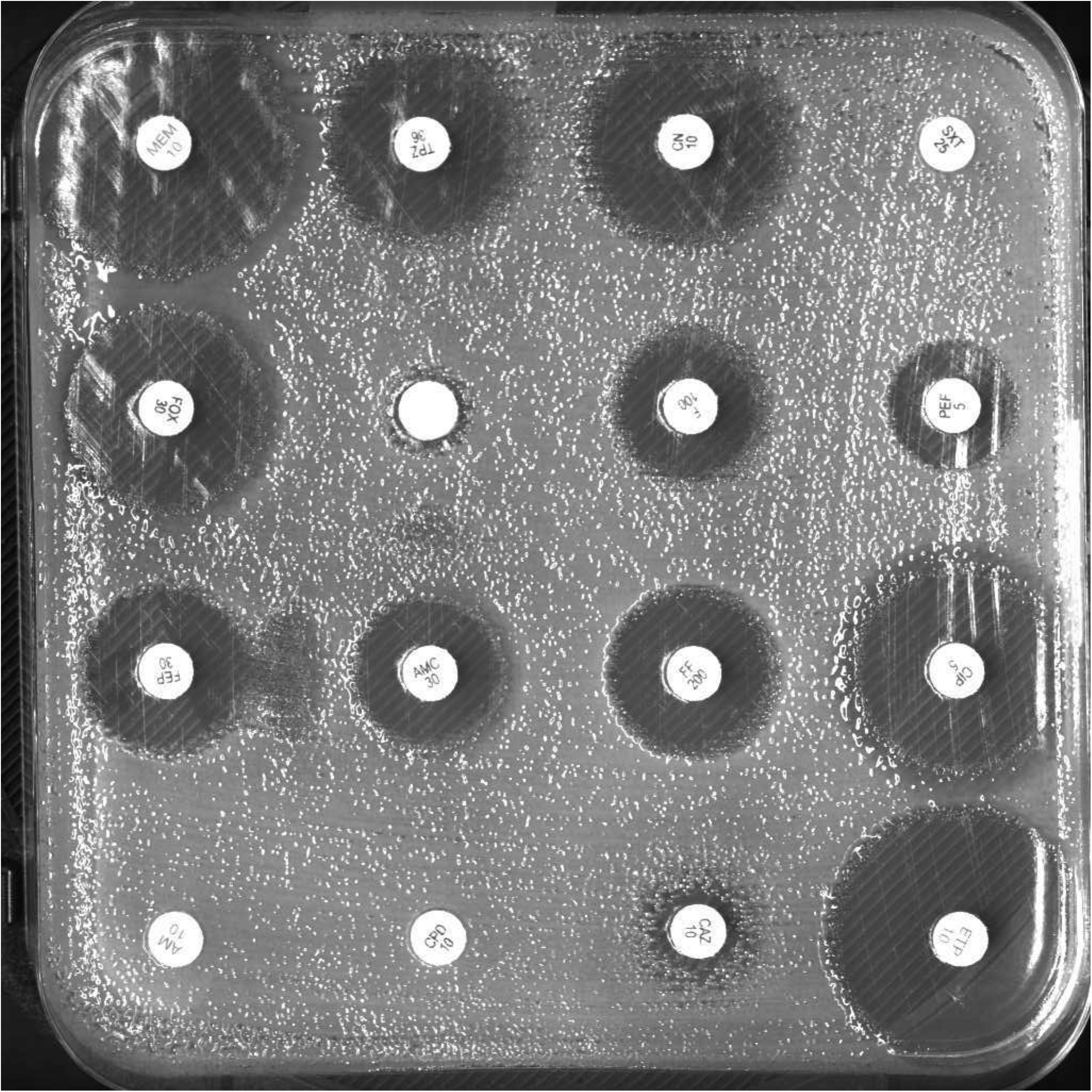
Representative photography of a disk diffusion assay generated by the SIRscan device. The image originates from routine diagnostics and shows no circles to indicate the measured inhibition zone diameter. As representative example image 2.3.1 was used.

**Supplementary Table 1.** A representative isolate (isolate 2.3.1) with measured inhibition zone diameters. The measured inhibition zones were interpreted according to EUCAST. As representative example image 2.3.1 was used. R, resistant; I, susceptible, increased exposure; S, susceptible; N. read, not read.

| Abbreviation | Antibiotic | Inhibition zone (mm) | Interpretation (S/I/R) | Resistant (mm) | Susceptible (mm) |
| --- | --- | --- | --- | --- | --- |
| CPD | Cefpodoxime | 6 | R | <21 | ≥21 |
| AMC | Amoxicillin/Clavulanic acid | 16 | R | <16 | ≥16 |
| CRO | Ceftriaxone | 6 | R | <22 | ≥25 |
| CIP | Ciprofloxacin | 23 | I | <22 | ≥25 |
| FOX | Cefoxitin | 22 | S | <19 | ≥19 |
| CAZ | Ceftazidime | 10 | R | <19 | ≥22 |
| TPZ | Piperacillin/Tazobactam | 19 | R | <20 | ≥20 |
| SXT | Sufamethoxazol-Trimethoprim | 6 | R | <11 | ≥14 |
| MEM | Meropenem | 30 | S | <16 | ≥22 |
| CN | Gentamicin (10ug) | 23 | S | <17 | ≥17 |
| F100 | Nitrofurantoin (100ug) | 15 | S | <11 | ≥11 |
| ETP | Ertapenem | 29 | S | <25 | ≥25 |
| FF | Fosfomycin | 17 | N. read | <24 | ≥24 |
| PEF | Pefloxacin | 13 | R | <24 | ≥24 |
| FEP | Cefepime | 18 | R | <24 | ≥27 |
| AM10 | Ampicillin (10ug) | 6 | R | <14 | ≥14 |

**Supplementary Table 2. Representative output table from the EUCAST-GPT-expert**. In this example sample 6.70.1: *Escherichia coli* is shown.

## Argumentation

- **None:** Based on the disk diffusion readings and adherence to EUCAST guidelines, there is no indication of reduced susceptibility that would suggest the presence of ESBL, AmpC, or carbapenemase resistance mechanisms. The inhibition zones are consistent with a susceptible Escherichia coli profile.
- **ESBL-production:** The inhibition zones for cefotaxime and ceftazidime are within the susceptible range without any signs of blunting, making ESBL production unlikely.
- **AmpC-production:** No diminished zone diameters for cefoxitin observed, suggesting AmpC production is unlikely. Furthermore, the susceptibility to amoxicillin-clavulanate supports the absence of AmpC.
- **Carbapenemase production:** The preserved zone sizes for meropenem and ertapenem indicate that carbapenemase production is very unlikely. There is no phenotypic evidence suggesting carbapenem resistance.

